# Growth defects and loss-of-function in synthetic gene circuits

**DOI:** 10.1101/623421

**Authors:** Evangelos-Marios Nikolados, Andrea Y. Weiße, Francesca Ceroni, Diego A. Oyarzún

## Abstract

Synthetic gene circuits perturb the physiology of their cellular host. The extra load on endogenous processes shifts the equilibrium of resource allocation in the host, leading to slow growth and reduced biosynthesis. Here we built integrated host-circuit models to quantify growth defects caused by synthetic gene circuits. Simulations reveal a complex relation between circuit output and cellular capacity for gene expression. For weak induction of heterologous genes, protein output can be increased at the expense of growth defects. Yet for stronger induction, cellular capacity reaches a tipping point, beyond which both gene expression and growth rate drop sharply. Extensive simulations across various growth conditions and large regions of the design space suggest that the critical capacity is a result of ribosomal scarcity. We studied the impact of growth defects on various gene circuits and transcriptional logic gates, which highlights the extent to which cellular burden can limit, shape and even break down circuit function. Our approach offers a comprehensive framework to assess the impact of host-circuit interactions *in silico*, with wide-ranging implications for the design and optimization of bacterial gene circuits.

---

Synthetic gene circuits rely on the machinery of the host cell where they reside. As the host fuels the expression of foreign genes, it diverts away resources from vital processes. This causes host cells to be susceptible to contextual effects known as “burden”, which may also affect the function of synthetic circuits^1,2^. The competition between synthetic and native genes produces a complex interplay between circuits and their host^3^. Such interplay perturbs endogenous processes and results in slow growth, reduced biosynthesis, and the onset of stress responses, all of which can affect circuit function^4,5^. Characterization of individual circuit parts is therefore insufficient for accurate prediction of function, and can thus lead to numerous rounds of trial-and-error iterations between circuit design, characterization and testing.

The seminal work by Tan and colleagues demonstrated that growth defects can dramatically change circuit function^6^. To improve our understanding of host-circuit interactions, a number of recent studies have focused on the cross-talk between gene circuits and host resources for transcription^7^, translation^2,8–11^ and protein degradation^12,13^. In particular, theoretical and experimental results showed that expression of heterologous proteins generally causes a decrease in constitutive expression of other genes^7–10^, due to limited resources for transcription and translation.

As gene circuits grow in size and complexity, their footprint on the host becomes a limiting factor on function. Ceroni and colleagues developed the first tool to experimentally quantify the burden caused by heterologous gene expression^8^. They built a “capacity monitor” consisting of a GFP-expressing cassette as proxy for the gene expression capacity in *Escherichia coli*. The GFP output of the monitor drops when heterologous gene expression is triggered, depending on the amount of resources sequestered by the synthetic construct. Subsequent works have focused on strategies to mitigate the deleterious impact of burden. For instance, Shopera *et al*^14^ found that negative feedback regulation reduces the cross-talk between gene circuits. Gorochowski *et al*^15^ showed that transcriptional profiles can be used to debug failure modes of various gene circuits. Most recently, Rugbjerg and colleagues^16^ coupled metabolite production to the expression of essential endogenous genes, which allowed to exploit evolutionary forces and increase production, while Darlington *et al* engineered a system that alleviates resource bottlenecks via an orthogonal ribosome^17^. Recent work further showed that promoters linked to the heat shock response of *E. coli* respond to heterologous expression, and then employed them to build a CRISPR/dCas9-based feedback system to control expression^18^.

Experimental quantification of growth defects, however, can be impractical because pushing a host into a burdened state may trigger stress responses and escape mutations^16,19^. This can shift the host physiology into new regimes that are strain-specific and generally not well understood. Together with appropriate experimental tools to better understand growth defects, there is growing need for predictive models that link circuit function to growth rate and the overall physiology of the host.

Our goal in this paper is to gain a quantitative grasp of the interplay between circuit function and growth defects. To this end, we use mathematical models for host-circuit systems that predict growth rate from the allocation of resources between native and foreign processes. We employ a mechanistic growth model^20^ extended with a cellular capacity monitor^8^ as a backbone for modelling gene circuits expressed in a bacterial host. Our models capture demands for cellular resources and their impact on endogenous genes for metabolism and other essential processes. The mechanistic nature of the models allows direct incorporation of tunable parameters that are commonly used in circuit design, such as the gene induction strength or ribosomal binding site (RBS) strength. Model simulations suggest that ribosomal scarcity can lead to growth defects and cause the capacity for gene expression to reach a tipping point. Such *critical capacity* appears to be a hallmark for the host transition into an overburdened state that impairs heterologous expression. We use extensive simulations to identify the implications of this phenomenon for circuit function and their corresponding design spaces.

## RESULTS AND DISCUSSION

### Mechanistic host-circuit models

We built several models for gene circuits coupled with the host physiology. We describe the host with a recently developed mechanistic growth model^20^, parameterized for proteome composition data in *E. coll*^21^. The growth model provides a dynamic description of how cells allocate their resources across various components of the proteome in midexponential growth phase. As illustrated in Fig. 1, we partition the proteome into four components: circuit proteins, ribosomes, metabolic enzymes, and housekeeping proteins. The model accounts for ribosomal autocatalysis, i.e. ribosomes require free ribosomes for their own expression, and accurately predicts specific growth rate emerging from the interplay between metabolism, biosynthesis and circuit activity. In the model, mRNA transcripts compete for free ribosomes and energy for translation. Heterologous circuit genes thus compete with native genes for cellular resources, while the predicted growth rate dilutes away the circuit proteins. This creates a two-way interaction between gene circuits and their cellular host. Our models can readily incorporate the impact of various growth media, as well as circuit design parameters such as gene induction, modelled as the maximal transcription rate of a gene, or RBS strengths, modelled via changes to the binding affinity between transcripts and ribosomes. Further details on model construction and parameterization can be found in the Methods.

**FIG. 1.**
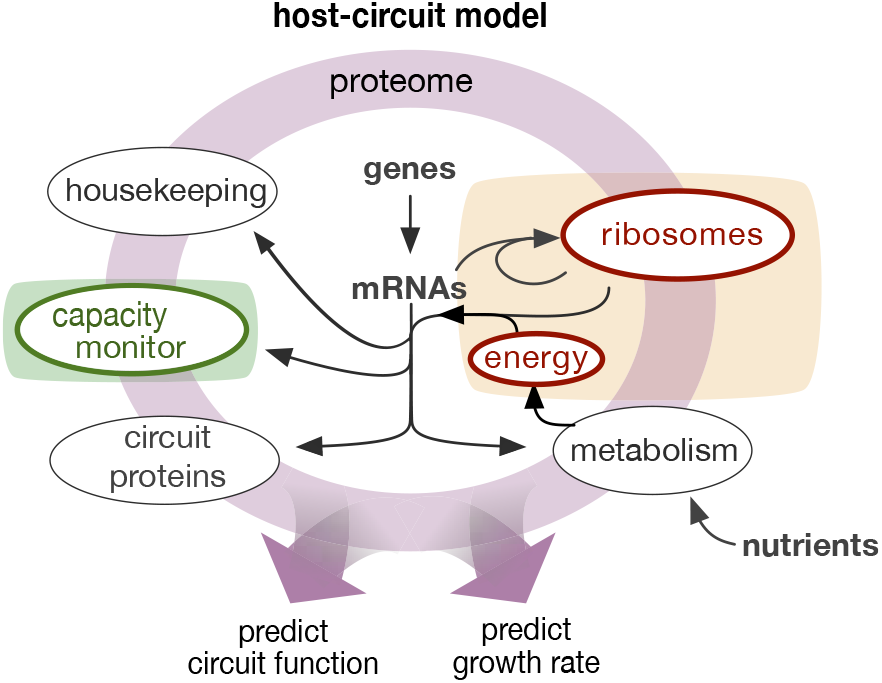
Integrated host-circuit model. We built mathematical models for gene circuits coupled with the physiology of a bacterial host. Circuit and native genes compete for free ribosomes and energy molecules. Our models are based on a mechanistic description of bacterial growth from a coarse-grained partition of the proteome^20^. The host model includes a GFP capacity monitor^8^ to track the impact of synthetic circuits on the cellular capacity for gene expression.

To assess the impact of the interplay between circuit and host, we conducted a series of simulations that mimic experiments commonly used in the design and optimization of circuits. Previous studies revealed trade-offs between gene induction and RBS strengths that shape cellular burden^8^. At the same time, different growth media have been reported to influence the ribosomal pool within cells^21^. To this end, we implemented various nutrient conditions, and explored the joint impact of RBS strength and circuit induction. All our simulations also include constitutive expression of a green fluo-rescent protein (GFP) gene. This corresponds to an *in silico* version of the “capacity monitor”^8^, where changes in constitutive GFP expression reflect competition for a shared pool of cellular resources. Here we define cellular capacity as the rate of GFP expression per cell^8^ in mid-exponential growth phase, which provides a convenient metric to quantify the impact of synthetic circuits on the resources of their host. In our models capacity corresponds to the steady-state rate of GFP expression.

### Limits on expression of heterologous proteins

We first focused on the expression of a heterologous protein from an inducible promoter for various gene induction strengths, RBS strengths and growth media. As shown in Fig. 2A, the model predicts that increased induction indeed causes an increase in heterologous expression. We found, however, that heterologous expression does not grow monotonically with induction, but instead protein output reaches a maximum and drops sharply for stronger gene induction. Simulations for different nutrient conditions and RBS strengths reveal that this phenomenon appears for a broad range of the induction-RBS design space, as well as in different growth conditions (Fig. 2B). Specifically, we note that the curves in Fig. 2A shift horizontally with RBS strength, which indicates that mRNA binding strength and circuit induction jointly increase expression output. At the same time, nutrient quality scales protein expression globally, with richer media resulting in higher protein output (Fig. 2B).

**FIG. 2.**
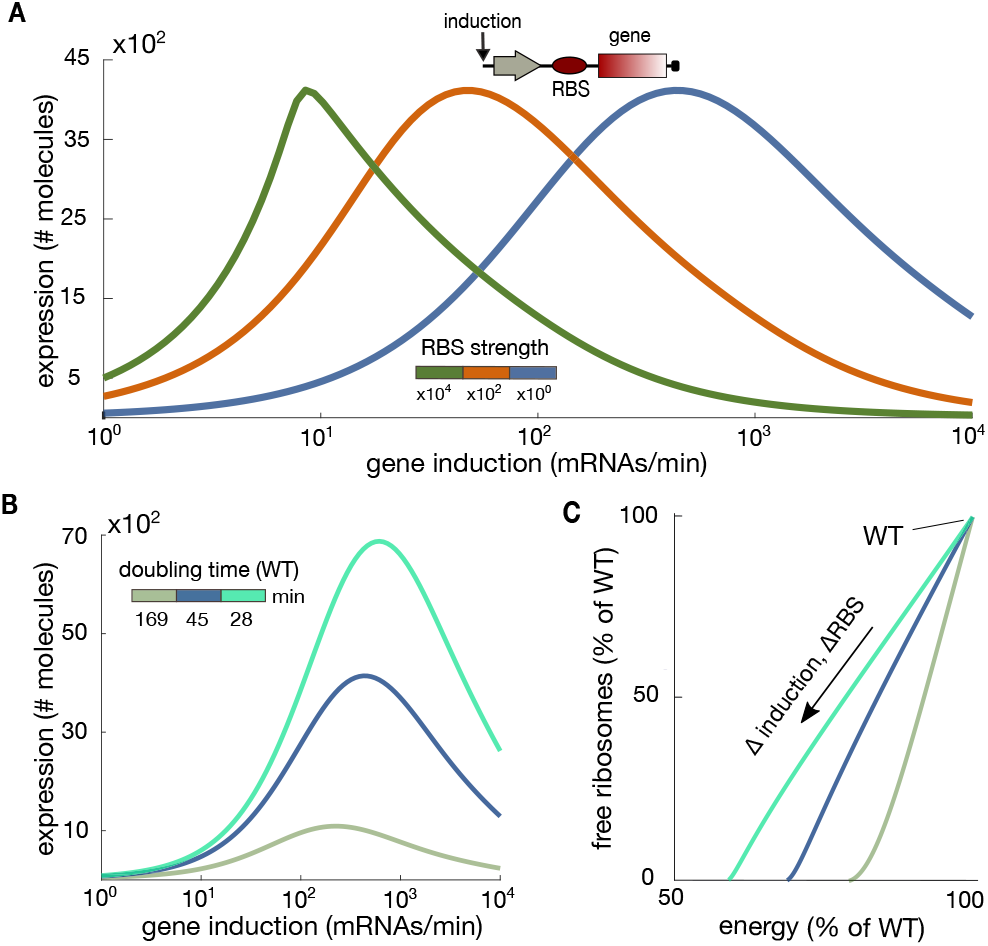
Limits on expression of an inducible gene. (**A**) Doseresponse curve of inducible gene expression for increasing induction and RBS strengths. The host-circuit model predicts a maximal expression output followed by a sharp decrease for strong induction. RBS strengths are fold-changes of the nominal case (in blue). (**B**) Expression of the inducible gene for various nutrient qualities; colours represent the predicted doubling time of the wild type strain. (**C**) Steady-state abundance of host resources (energy and free ribosomes) for increasing gene induction strength; colors correspond to the legend in panel B. The energy-ribosome relation is independent of the heterologous RBS strength. The wild type (WT) corresponds to the case of nil induction. Model predictions are in mid-exponential growth phase and parameters can be found in Methods C.

The observed impact of RBS strength and nutrient quality (Fig. 2A–B) suggest that limits in heterologous expression may be linked to energy and ribosome availability. We thus examined the availability of energy molecules and free ribo somes predicted by the model (Fig. 2C). In all cases, we observed a depletion of free ribosomes for high circuit induction, while energy levels remained above 60% of the wild type. Richer growth media result in more energy consumption, yet we found that the relation between energy and free ribosomes is unaffected by the RBS strength. To explain this phenomenon, from the model we derived a relation for the steady-state levels of energy molecules:

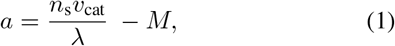

where *a* represents abundance of energy molecules, *n_s_* is the nutrient quality, *v*_cat_ is the rate of nutrient metabolization, *M* is the cell mass, and λ is the predicted growth rate (details in Methods). The formula in (1) indeed does not depend on the RBS strength of the heterologous gene, in agreement with Fig. 2C.

### Critical cellular capacity

To better characterize the limits of heterologous expression, we examined the dependency of cellular capacity (quantified by the GFP capacity monitor^8^) and growth rate on the gene induction strength. Across various growth conditions, we found two regimes where circuit and host display qualitatively distinct behavior. As shown in Fig. 3A, for weak to moderate induction, there is a near-linear dependency between cellular capacity and circuit output. In this regime, increased induction leads to a larger protein output at the expense of a decrease in cellular capacity. This is in agreement with previous experiments in which expression of an inducible gene caused a drop in constitutive expression of another gene^7,9,10^. The lowered cellular capacity is accompanied by a growth defect of up to “~50%. with respect to the wild type strain (see Fig. 3B).

**FIG. 3.**
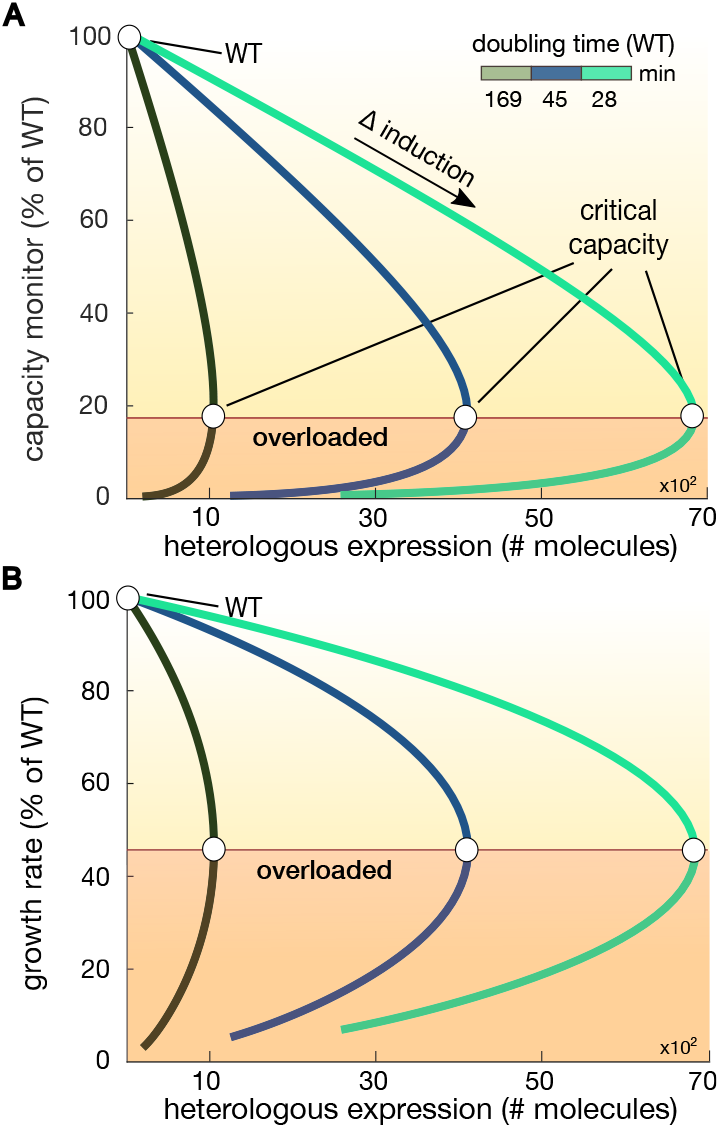
Critical cellular capacity and growth defects. **(A)** Output of the capacity monitor^8^ with respect to expression of the inducible gene in Fig. 2A. We found a critical capacity at which the host transitions into an overloaded regime. In the overloaded regime, both inducible expression and growth rate decrease with the induction strength. Nutrient quality shifts the capacity curves to higher expression levels but does not alleviate the resource bottlenecks. **(B)** At the tipping point, an ~80% drop in capacity is accompanied by growth defects of ~50% for the considered *E. coli* model strain. The wild type (WT) corresponds to the case of nil induction. Model predictions are in mid-exponential growth phase and parameters can be found in Methods C.

Beyond the above regime, however, our simulations reveal a tipping point at which the relation between growth and circuit output changes drastically. As seen in Fig. 3A, in this new regime, both cellular capacity and protein output jointly decrease. The tipping point between both regimes corresponds to a *critical cellular capacity*, which is associated with a depletion of available ribosomes observed in Fig. 2C. Beyond the tipping point, the host enters an overloaded regime in which scarcity of ribosomes is forbidding for the expression of both endogenous and heterologous genes.

We further employed the host-circuit models to explore how induction and RBS strength jointly impact cellular growth and protein expression. As shown in Fig. 4A, for similar levels of induction, designs with stronger RBS impart more burden to the host. The relation between growth rate and induction strength becomes more sensitive for stronger RBS. In particular, for designs with a ~100-fold increase in RBS strength, small changes in gene induction lead to ~60% change in growth rate. This is a form of growth ultrasensitivity that emerges as a result of excessive burden on the host. Simulations also indicate that heterologus expression is limited even for increased RBS strength (Fig. 4B). We also found that the whole design space, i.e. all combinations of induction and RBS strength, collapse to a single curve of expression vs. growth (Fig. 4C), where the tipping point in capacity and its corresponding drop in growth rate become apparent.

**FIG. 4.**
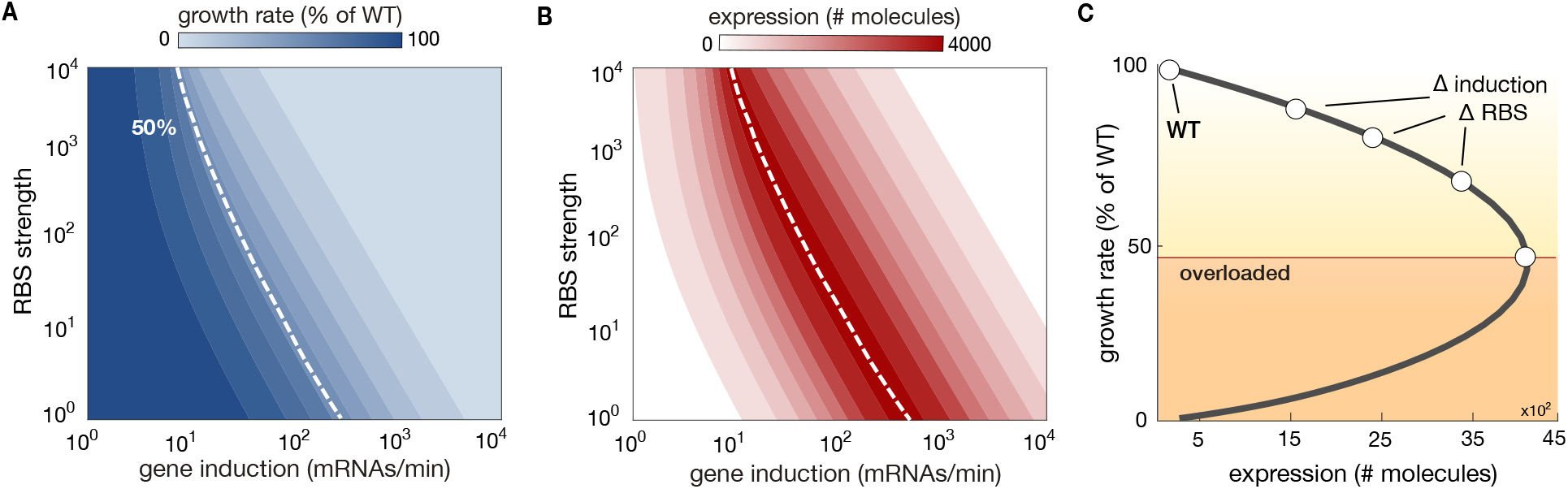
Design space for the expression of an inducible gene. **(A)-(B)** Predicted growth rate and heterologous expression upon changes in gene induction and RBS strength for the inducible gene in Fig. 2A. A stronger RBS shifts the critical capacity (white, dashed line) to weaker induction values. For strong RBS, growth rate becomes more sensitive to changes in gene induction. **(C)** The whole parameter space collapses to a single curve similar to Fig. 3B. Wild type (WT) corresponds to the case of nil induction.

### Loss-of-function in gene circuits

To explore the relation between growth defects and circuit function, we studied the dynamics of several gene circuits. Using our host-circuit models, we systematically investigated their function and impact on growth rate across large ranges of their design parameters.

#### Genetic toggle switch

We first investigated the effects of RBS strength on the design space of the toggle switch^22^ (Fig. 5A). This circuit displays bistability and can be toggled between two steady states with external inducers. The design space of the toggle thus contains all combinations of inducer levels that lead to a bistable system. Our simulations (Fig. 5A) predict that stronger RBS causes a substantial reduction of the design space, as a result of growth defects. Such shrinkage is not observed in isolated models of the switch, suggesting that competition for host resources causes a reduced design space and loss-of-function. The hysteresis diagrams in Fig. 5B show that within the bistable regime, the switching dynamics are also affected by strong RBS, causing the “high” state to decrease monotonically upon stronger induction.

**FIG. 5.**
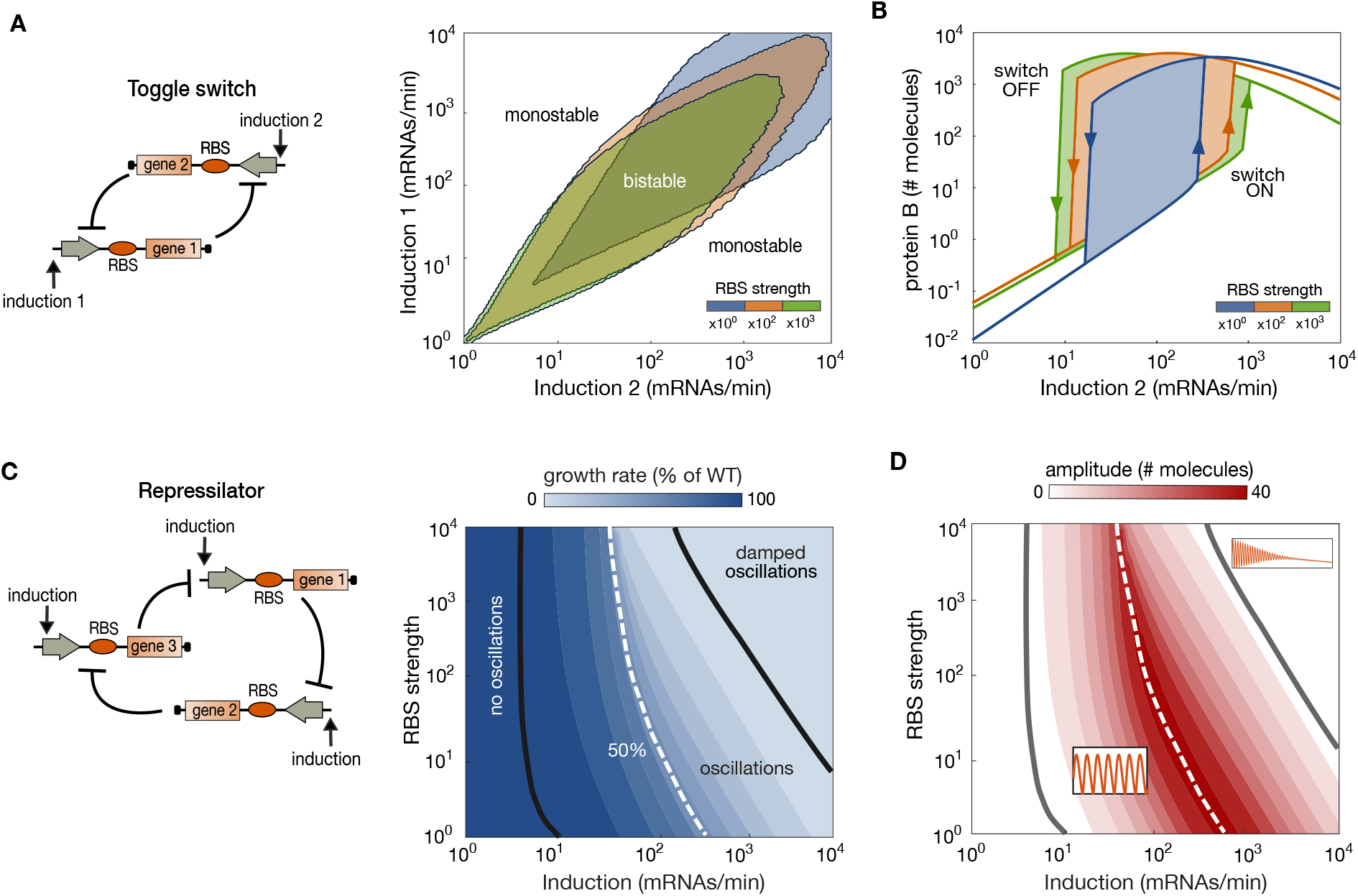
Impact of host-circuit interactions on a genetic toggle switch and a genetic oscillator. **(A)** Circuit diagram of the genetic toggle switch^22^ and predicted regions for bistability for varying inductions of both genes. Our host-circuit model predicts a shrinkage of the design space for stronger RBS. **(B)** Hysteresis diagrams of the toggle switch reveal changes in circuit function due to host-circuit interactions; the expression level of the ON state drops for strong induction of one gene. Hysteresis diagrams were computed by fixing the induction level of gene 1 and varying the induction of gene 1. **(C)** Circuit diagram of the repressilator^23^ and predicted growth rate across the design space. Host-circuit models predict a decrease in growth rate for increasing induction and RBS strength of the repressilator. The white dashed line represents designs with a 50% drop in growth rate with respect to the wild type (WT). The WT model corresponds to the host model coupled with the GFP capacity monitor only. **(D)** Similar to the toggle switch in panel A, the host-circuit model predicts a shrinkage of the design space for oscillations for strong induction and RBS; insets show the circuit dynamics in each regime. Details on model parameters and simulations can be found in Methods.

#### Genetic oscillator

Next, we explored the impact of design parameters on the repressilator^23^, one the most well studied genetic oscillators (Fig. 5C). Upon changes in induction and RBS strengths, we found a monotonic drop in growth rate, similarly as in the expression of the inducible promoter in Fig. 4A. We found a large design space for oscillations, covering several orders of magnitude in induction and RBS strengths. Outside the design space, we found two additional regimes: designs that display a single stable steady state and thus lack oscillations, and designs that display damped oscillations that decay to a steady state. The first regime is consistent with what is observed in the isolated circuit^23^, but the damped oscillations appear to be an emergent feature from the cross-talk between circuit and the host.

As seen in Fig. 5D, changes in growth are accompanied by variations in the amplitude of oscillations, which reach a maximal value for ~50% drop in growth rate. For stronger RBS and gene induction, we observe a sharp drop in the oscillation amplitude, until complete loss-of-function. This is not seen in isolated circuit models and thus reflects the impact of host physiology on the design space and dynamics of the oscillator. Akin to our analysis of heterologous expression and the toggle switch, our simulations (Fig. 5C–D) suggest that the repressilator is subject to a critical cellular capacity that defines a transition from normal operating condition to a high-burden, slow growth regime. The critical capacity appears within the oscillation-positive region suggesting an additional limitation for the available design space.

#### Genetic logic gates

To illustrate the impact of host-circuit interactions on more complex genetic circuitry, we examined several logic gates based on transcriptional regulators^24^. We simulated NOT, AND and NAND gates (Fig. 6A, C, E) and predicted the resulting growth rates for combinations of inputs (Fig. 6B, D, F). As seen in Fig. 6B (left), the function of the NOT gate remains largely unaffected by host-circuit interactions, even for strong RBS of gene 2. For intermediate input levels, however, simulations in Fig. 6B (right) predict an increase in growth rate of up to 40% with respect to the basal case; this effect is more pronounced for designs with weaker RBS of gene 2. Such apparent growth benefit is a consequence of the circuit architecture (Fig. 6A): an increase in the input causes a stronger repression of gene 2 and thus relieves the burden on the host. But since the expression of the repressor coded by gene 1 also causes burden, for sufficiently high inputs the expression of gene 1 counteracts the growth advantages gained by repression of gene 2, resulting in an overall drop in growth rate.

**FIG. 6.**
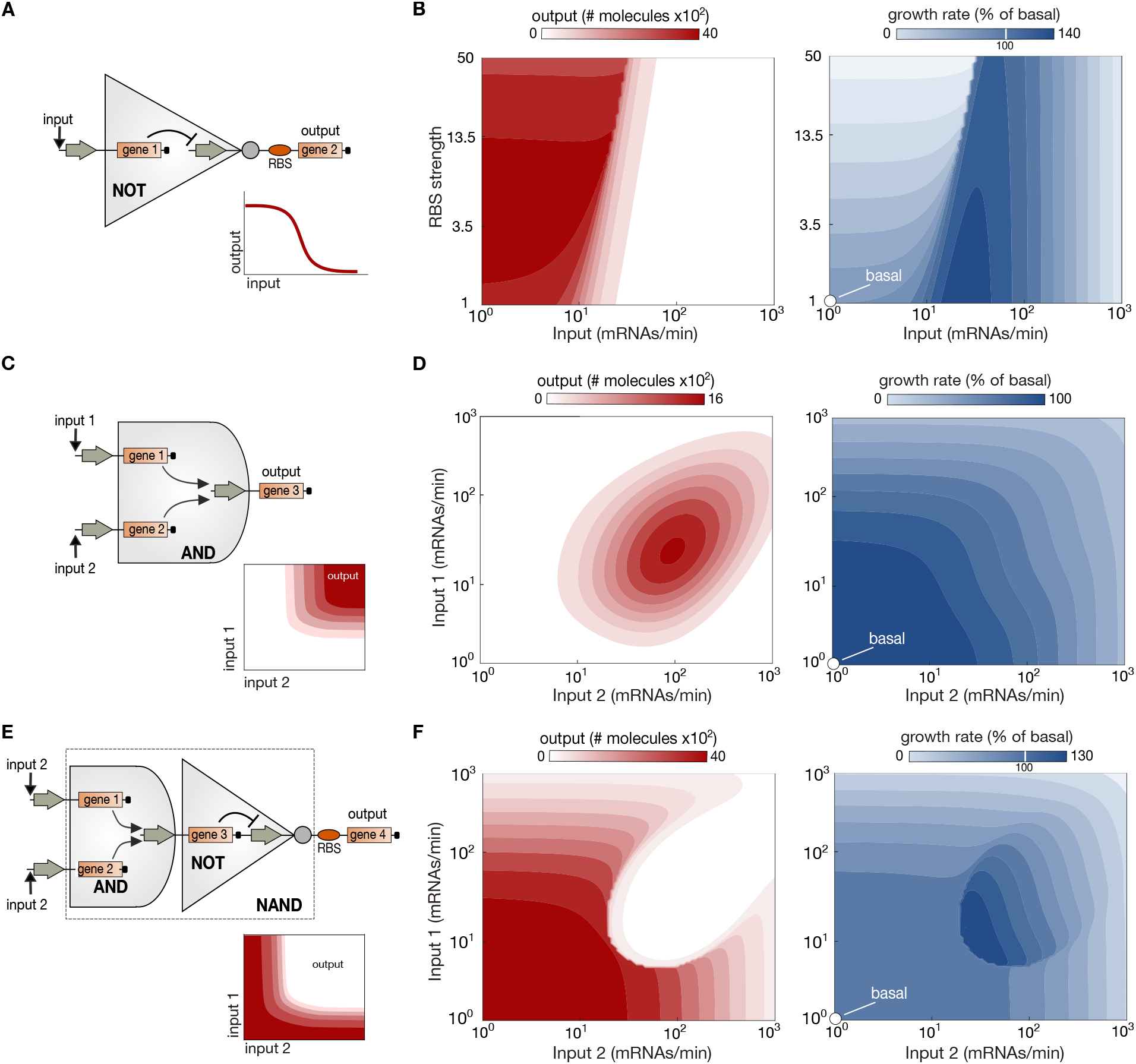
Impact of host-circuit interactions on genetic logic gates. **(A)** Schematic of a NOT gate and its ideal response curve. **(B)** Predicted response of NOT gate and growth rate for increasing RBS strength of gene 2. **(C)** Schematic of an AND gate and its ideal response surface. **(D)** Predicted response of AND gate and growth rate of the host. **(E)** Schematic of a NAND gate and its ideal response surface; the gate is the composition of AND and NOT gates from panels A and C. **(F)** Predicted response of a NAND gate and growth rate of the host. All gates are based on an implementation using transcriptional activators and repressors^24^; model details and parameters can be found in the Methods.

In contrast, simulations of the AND gate in Fig. 6D (left) suggest a much stronger impact of host-circuit interactions. An ideal AND gate should produce a high output when both inputs are high, see Fig. 6C (inset), yet the host-circuit models predict a bell-shaped response surface, where the output reaches a maximal value for an intermediate level of the inputs, beyond which the output drops monotonically. The loss-of-function coincides with a drop in growth rate observed for increased levels of either input, as seen in Fig. 6D (right).

The composition of both circuits, the NAND gate in Fig. 6E, displays a complex relation between inputs and output. The gate produces appropriate responses in large ranges of the input space (Fig. 6E, left), with some distortions possibly caused by the loss-of-function of the AND component. Likewise, we observe a growth advantage for intermediate levels of the input as a result of the architecture of the NOT gate, akin to what we observed in Fig. 6B (right).

## DISCUSSION

Microbes tailor their proteome composition to sustain growth and other vital functions. Heterologous expression perturbs the homeostatic balance of a cellular host by drawing resources for biosynthesis, which in turn can lead to unpredictable circuit behaviour^2^. Such multi-faceted interactions between circuits and their host are difficult to predict, yet they are thought to be major contributors to poor functionality in synthetic gene circuits^1^.

In this work we employed mechanistic host-circuit models to study the interplay between circuit function and growth defects. The models consistently predict the existence of a critical capacity for gene expression. Stronger gene induction results in higher heterologous expression, but only up to a maximal yield achieved at the critical capacity and with a substantial growth defect. The critical capacity defines a state in which heterologous expression becomes severely hampered by the innate resource limitations in the host. If gene induction is strong enough to tip the host over the critical capacity, the relation between induction and protein yield reverses, with stronger induction causing reduced yield in heterologous expression and a further decrease in growth rate. The critical capacity thus provides a quantitative metric to quantify how a host transitions into a burdened regime limited by ribosomal availability.

Our models suggest that the main source of burden is the depletion of free ribosomes for translation of circuit genes, in agreement the widespread conception that ribosomal availability is a major control node for cellular physiology^21,25,26^. Also in agreement with recent experiments^8,9^, the host-circuit models indicate that designs with stronger RBS impose a heavier burden on the host. An increased mRNA-ribosome binding affinity gives foreign transcripts a binding advantage, resulting in increased burden and a smaller pool of free ribosomes available for native transcripts. Thanks to the unique ability of our models to simultaneously predict circuit function and specific growth rate, we can map cellular burden into quantitative predictions for growth defects.

We also found that host-circuit interactions can drastically change the function of synthetic constructs. We first focused on the genetic toggle switch^22^ and the repressilator^23^, two exemplar circuits designed to achieve specific dynamic responses. In both cases we found that the circuit design spaces shrink as a result of the interplay with the host physiology. For the toggle switch, strong RBS leads to loss of function for gene inductions that would otherwise be predicted as producing a bistable response. A similar phenomenon appeared in the repressilator, where strong RBS may cause the circuit to lose the oscillatory function. Simulations of genetic logic gates^24^ further illustrate the impact of host-circuit interactions, suggesting that burden may pervade the function of a wide range of genetic circuitry.

A number of studies have proposed models for resource competition and its impact on the function of gene circuits^2^. The concept of “retroactivity”, introduced by del Vecchio and colleagues^27^ provides a useful metric to analyze how circuits affect each other’s function due to limited resources^13^. Other approaches have focused on minimal models to describe competition for a finite pool of resources^7,9,10^. Important limitations of these previous works are the assumption of a constant, circuit-independent, growth rate and the lack of explicit mechanisms that link circuit expression to the abundance of free ribosomes. In regimes where growth defects are small, our host-circuit models agree with previous works^7–10^ that show a decrease in constitutive expression in response to the induction of heterologous genes. These observations are in agreement with the near-linear relation we found between the expression of inducible and constitutive genes before reaching the critical capacity (Fig. 3A). However, the ability of our models to predict growth rate and shifts in proteome composition offers additional information on how circuits behave within their cellular context. The mechanistic nature of the models allows us to directly simulate the impact of design parameters such as RBS strengths, gene induction and protein degradation tags. Moreover, because the models include ribosomal autocatalysis, they account for cases in which strong RBS leads to a sizeable fraction of the ribosome pool being sequestered by heterologous transcripts, which amplifies the impact of burden by limiting the ability of the host to synthesize more ribosomes and alleviate the resource bottleneck. Our mathematical models also account for various growth media and their impact on the host-circuit interplay. In particular, simulations suggest that richer media do not mitigate the resource bottleneck, but instead allow for increased expression of both endogenous and heterologous genes.

In this paper we focused on host-circuit competition for energy and free ribosomes, two key cellular resources that impact growth directly. In practice, gene circuits also consume other cellular components that may become resource bottlenecks, such as RNA polymerases and *σ*-factors for transcription, or amino acids and tRNAs for translation. Some of these components can be readily included in mechanistic models similar to ours^28^, yet this can increase model complexity and obscure the relations between different sources of burden. Recent experimental work^29^ has shown that engineered metabolic pathways can impose an additional source of burden resulting from metabolic imbalances. This offers exciting avenues for modelling the activity of engineered pathways coupled with the host physiology, with promising applications at the interface of metabolic engineering, synthetic biology and mechanistic cell modelling^30^.

Experimental validation of our predictions would greatly help the identification of the critical capacity barrier *in vivo*. Recent datasets suggest highly nonlinear relations between growth rate and heterologous expression^31^, raising compelling prospects for the integration of mechanistic cell models with such large-scale characterization data. Moreover, since burden can also lead to the insurgence of escape mutants^8,16,19^, future experimental work may focus on characterizing the correlation between critical capacity and loss of circuit function due to mutations. Our mechanistic models hold promise as detailed and tractable descriptions of host-circuit interactions. Their level of granularity finds an adequate compromise between simple phenomenological models for growth^21^ and complex models for whole-cell physiology^32^. This modelling framework thus offers a powerful quantitative tool for teasing out the interplay between burden and circuit function, ultimately, paving the way for more robust and predictable synthetic biology.

## METHODS

### A. Model for a bacterial host

We use the coarse-grained model described in Weiße et al.^20^. The model accounts for the uptake of an external nutrient at a fixed concentration *s* by a transport protein *p_t_*. The internalised nutrient *s_int_*, is catabolised by a metabolic enzyme *p_m_* to produce a generic form of energy, denoted *a*, that models the total pool of intracellular molecules required to fuel biosynthesis, such as ATP and aminoacids. The model includes transcription and degradation of mRNAs *m_x_*, their binding to free ribosomes *p_r_* to form ribosome-mRNA complexes *c_x_*, and translation reactions for synthesis of the various components of the proteome *p_x_*. We divide the components of the proteome (*p*_x_) into several classes *x* ∈ {*r, t, m, q*, gfp}. The *q* class represents house-keeping proteins and gfp corresponds to the “capacity monitor” developed by Ceroni and colleagues^8^. The model reactions are displayed in Table I.

**TABLE I.** Reactions of the wild type host model. The proteome components are *x* ∈ {*r, t, m, q*, gfp}.

| metabolism | transcription | ribosome binding |
| --- | --- | --- |
| $s \xrightarrow{v_{\text{imp}}} s_{\text{int}} \xrightarrow{v_{\text{cat}}} n_s a$ | $\emptyset \xrightarrow{w_x} m_x$ | $p_r + m_x \xrightleftharpoons[k_u]{k_b} c_x$ |
| degradation | translation | dilution |
| $m_x \xrightarrow{d_{m,x}} \emptyset$<br>$p_x \xrightarrow{d_x} \emptyset$ | $n_x a + c_x \xrightarrow{v_x} p_r + m_x + p_x$ | $a, m_x, c_x, p_x \xrightarrow{\lambda} \emptyset$ |

In the model, λ is the predicted growth rate and the nutrient efficiency parameter *n_s_* determines energy yield per molecule of internalized nutrient. The model includes dilution of all chemical species by cell growth, as well as degradation of transcripts and proteins (with rate constants *d*_m,x_ and *d*_x_, respectively). The specific growth rate can be computed from the total number of ribosomes engaged in translation^20^:

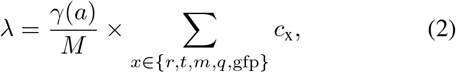

where *γ*(*a*) is the rate of translational elongation and assumed be a saturable function of the available energy, *γ*(*a*) = *γ*_max_*a*/(*a* + *K_γ_*) and *M* is the cell mass.

#### Proteome components

From the chemical reactions in Table I, we can write an ordinary differential equation (ODE) model for each component of the proteome:

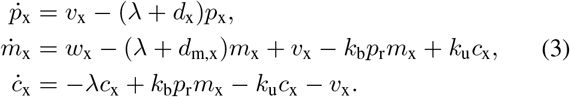

where *x* includes all proteins except ribosomes, i.e. *x* ∈ {*t, m, q*, gfp}. The equations for ribosomes are different because of ribosomal autocatalysis and competition for free ribosomes (see next section). Note that in Eq. (3) each gene requires three species: the transcript (*m*_x_), the mRNA-ribosome complex (*c*_x_), and the protein itself (*p*_x_). We assume that transcription and translation rates (*w*_x_ and *v*_x_, resp.) depend on the energy resource, *a*, and follow *w*_x_ = *w*_xmax_*a*/(*θ*_x_ + *a*) for all proteins except the housekeeping proteins, i.e. *x* ∈ {*r, t, m*, gfp}. The transcription of housekeeping mRNAs is subject to negative autoregulation so as to keep constant expression levels in various growth conditions, i.e. 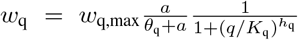. The translation rate for all proteins follows *v*_x_ = (*c*_x_/*n*_x_) × (*γ*_max_*a*/(*a* + *K_γ_*)) with *x* ∈ {*r, t, m, q*, gfp}, with *n*_x_ being the protein length in number of amino acids.

#### Shared cellular resources

In our model genes compete for two resources, an energy resource (*a*) and free ribosomes for translation (*p*_r_). Energy is obtained from the internalized nutrient, which we model by the ODE:

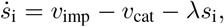

where nutrient import and catabolism are assumed to follow Michaelis-Menten kinetics of the form *v*_imp_ = *p*_t_*v*_t_*s*/(*K*_t_ + *s*) and *v*_cat_ = *p*_m_*v*_m_*s*_i_/(*K*_m_ + *s*_i_), with kinetic parameters *v*_t_, *K*_t_, *v*_m_ and *K*_m_. The ODEs for energy, free ribosomes, ribosomal transcripts and their complexes are:

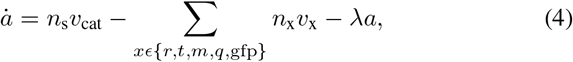

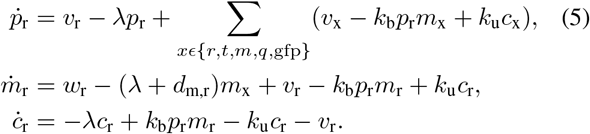

Eq. (5) accounts for ribosomal autocatalysis, i.e. ribosomal transcripts sequester free ribosomes for their own translation, and the pool of free ribosomes can increase both due to translation of new ribosomes and the free-up of ribosomes engaged in translation.

### B. Integrated host-circuit models

To model the expression of heterologous genes, we used equations analogous to Eq. (3) for each circuit gene:

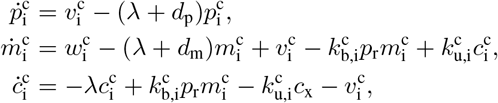

where the superscript c represents a circuit species and the subscript *i* denotes the *i*^th^ circuit gene. To model the impact of gene circuits in host resources, we modified the resource equations in (4)–(5):

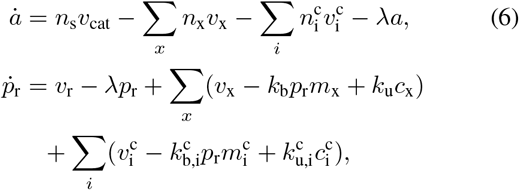

and the specific growth rate in Eq. (2) now includes translation of circuit genes:

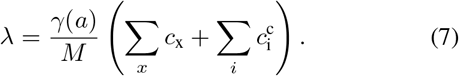

We modelled the translation rate of all foreign genes similarly as that of native genes:

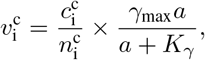

with 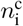 being the length of the *i*^th^ circuit protein. Finally, the transcription rate for foreign genes is:

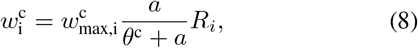

where 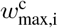 is the maximal transcription rate, and *R_i_* is a regulatory function for each circuit. For the inducible gene (Fig. 2–4) we use *R*_1_ = 1. The genetic switch (Fig. 5A–B) has two genes with regulatory functions:

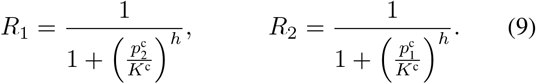

The repressilator (Fig. 5C–D) has three genes with regulatory functions:

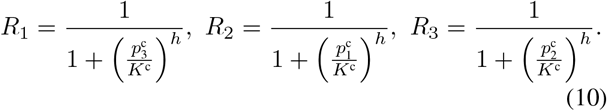

The NOT gate (Fig. 6A–B) has two genes with regulatory functions:

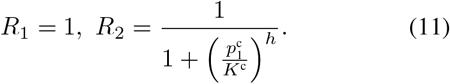

The AND gate (Fig. 6C–D) has three genes with regulatory functions:

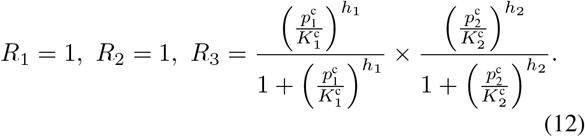

Finally, the NAND gate (Fig. 6E–F) has four genes with regulatory functions:

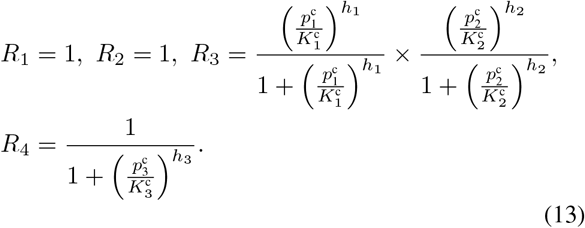

In the regulatory functions in Eq. (9)–(13) the parameters *K*^c^ and *h* represent a regulatory threshold and effective Hill coefficient, respectively.

### C. Model parameters

#### Model for host

We parameterized the wild type host model using parameters from Weiße *et al*^20^, which were estimated using Bayesian inference on *E. coli* growth data^21^. Throughout this paper we consider the wild type strain to already include the GFP capacity monitor from Ceroni et al^8^. The parameters for the capacity monitor are *w*_GFP,max_ = 100 mRNAs/min for the maximal transcription rate, and RBS equal to that of endogenous transcripts RBS_GFP_ = *k*_b_/*k*_u_ = 0.95 × 10^−2^ molecules^−1^. The transcripts and protein halflives of the capacity monitor were assumed to be two and four minutes, respectively, so that *d*_m,gfp_ = ln 2/2 and *d*_gfp_ = ln 2/4. We assume all other proteins are not subject to active degradation, i.e. *d*_x_ = 0 for *x* = {*r, m, t, q*}.

**TABLE II.** Model parameters for the host. All values taken from Weiße et al^20^; the mRNA-ribosome binding rate which was set to *k*_b_ = 0.95 ×10^−2^ min^−1^ molecules^−1^. Units of aa correspond to number of amino acids per cell.

| Name | Value | Name | Values |
| --- | --- | --- | --- |
| $s$ | $10^4$ (molecules) | $M$ | $10^8$ (aa) |
| $n_r$ | 7459 (aa/molecules) | $\theta_r$ | 427 (molecules) |
| $\gamma_{\text{max}}$ | 1260 (aa/min molecules) | $K_\gamma$ | 7 (molecules) |
| $v_t$ | 726 (min<sup>-1</sup>) | $K_t$ | 1000 (molecules) |
| $v_m$ | 5800 (min<sup>-1</sup>) | $K_m$ | 1000 (molecules) |
| $w_{r,\text{max}}$ | 930 (molecules / min) | $w_{m,\text{max}}, w_{t,\text{max}}$ | 4.14 (molecules / min) |
| $w_{q,\text{max}}$ | 949 (molecules/min) | $d_{m,r}, d_{m,t}, d_{m,m}, d_{m,q}$ | 0.1 (min<sup>-1</sup>) |
| $K_q$ | 152219 (molecules) | $h_q$ | 4 |
| $\theta_q, \theta_t, \theta_m, \theta_{\text{gfp}}$ | 4.38 (molecules) | $k_u$ | 1 (min<sup>-1</sup>) |
| $n_q, n_t, n_m, n_{\text{gfp}}$ | 300 (aa/molecules) | $n_r$ | 7459 (aa/molecules) |

#### Models for gene circuits

For all heterologous genes, we assumed transcript and protein half-lives of two and four minutes, respectively^23^, so that *d*_m_ = ln2/2 and *d*_p_ = ln2/4. We set the transcriptional energy threshold equal to that of native non-ribosomal genes, with *θ*^c^ = *θ*_x_ = 4.38 molecules for *x* ∈ {*q, t, m*, gfp}. Heterologous proteins were assumed to be of length 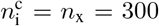 amino acids, equal to that of native non-ribosomal proteins.

The parameters for the gene circuits were chosen so as to produce realistic molecule numbers across all cases. We fixed the parameters of the toggle switch in Eq. (9) and re-pressilator in Eq. (10) to *K*^c^ = 100 molecules and Hill coefficient *h* = 2. For the NOT gate, we fixed the parameters of *R*_2_ in Eq. (11) to *K*^c^ = 250 molecules and h = 2. For the AND gate, we fixed the parameters of R_3_in Eq. (12) to 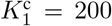 molecules and *h*_1_ = 2.381 for the activation by gene 1, and we used 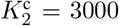 molecules and *h*_2_ = 1.835 for the activation by gene 2, similar to the parameter values estimated in Wang *et al*^24^. We modelled the NAND as the composition of the AND and NOT gate with RBS for gene 3 set to unity, without changing their individual parameters. The remaining circuit parameters were varied across simulations, as explained next.

### D. Circuit simulations

All simulations were computed with the stiff ODE integration routine `ode15s` in Matlab R2018a. For each circuit simulation, we initialize the native species of the host from numerical steady states computed from a separate simulation of the wild type strain. We modelled gene induction via the maximal transcription rate 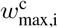 in Eq. (8). We modelled changes to RBS strength by varying the mRNA-ribosome binding rate constant 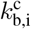 (in units of min^−1^ molecules^−1^) and the dissociation rate constant 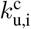 (in units of min^−1^). The quality of the growth media was modelled via the nutrient efficiency *n*_s_. Next we detail the parameter ranges for all simulations.

#### Inducible gene

In Fig. 2A-C we swept the gene induction in the range 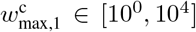. The RBS strengths in Fig. 2A were simulated with pairwise ratios between binding rate constants 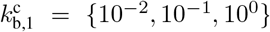 and dissociation rate constants 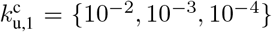; nutrient quality in Fig. 2A was fixed to *n_s_* = 0.5. In Fig. 2B–C we fixed RBS strength using 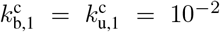 and varied the nutrient quality *n_s_* = {0.1,0.5,1}. In Fig. 3 we swept 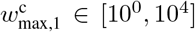 with nutrient quality *n_s_* = {0.1,0.5,1}. In Fig. 4 we swept 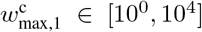 with fixed nutrient quality *n_s_* = 0.5. For the RBS strength we simultaneously increased the binding rate constant from 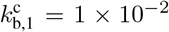 to 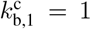 and decreased the dissociation rate constant from 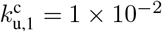 to 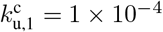.

#### Genetic toggle switch

To compute the regions for bistability in Fig. 5A we swept induction of both genes in the range 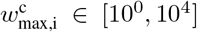, and numerically computed the number of stable steady states from a large number of model simulations starting from different initial conditions. To compute the hysteresis diagrams in Fig. 5B, we fixed the induction of protein 1, 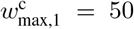, and varied 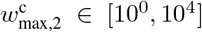 with RBS strengths using 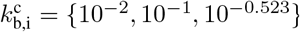, and 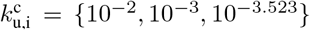 for both genes. The nutrient quality was fixed to *n_s_* = 0.5 in all toggle switch simulations.

#### Genetic oscillator

In Fig. 5C-D we swept the gene induction in the range 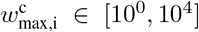. We changed the RBS strength by increasing the binding rate constant from 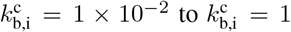, while simultaneously decreasing the dissociation rate constant from 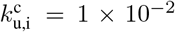 to 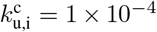 for the three genes; nutrient quality was fixed to *n_s_* = 0.5. Oscillations and their amplitude were detected using custom Matlab scripts.

#### Genetic logic gates

In all gates we fixed the nutrient quality to *n_s_* = 0. 5 and modelled the inputs via the maximal transcription rate 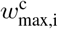 in the range 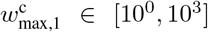. For the NOT we changed RBS strength of gene 2 by increasing the binding rate constant from 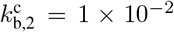 to 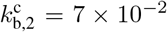, while decreasing the dissociation rate constant from 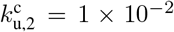 to 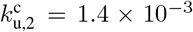. For the AND gate and NAND gates we fixed the RBS strength with 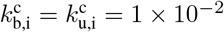 for all genes.

### E. Relation between energy and free ribosomes

As shown in Fig. 2C, model simulations suggest that the relation between energy and free ribosomes for various induction strengths is independent of the RBS strength of a heterologous gene. To explain this phenomenon, we can solve for *a* in (6) with *i* = 1 (inducible gene) at steady state:

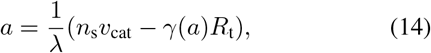

where the term 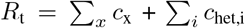 is the total number of ribosomes engaged in translation. From the formula for growth rate in (7) we get *γ*(*a*)*R_t_* = λ*M*, and after substitution in Eq. (14) we get the expression in Eq. (1) of the main text.

## AUTHOR CONTRIBUTIONS

E.-M.N. built the mathematical models and simulation code. All authors contributed to analysis of the results and writing of the manuscript.

